# The effect of the size of gold nanoparticle contrast agents on CT imaging of the gastrointestinal tract and inflammatory bowel disease

**DOI:** 10.1101/2024.01.20.576354

**Authors:** Derick N. Rosario-Berríos, Amanda Pang, Leening P. Liu, Portia S. N. Maidment, Johoon Kim, Seokyoung Yoon, Lenitza M. Nieves, Katherine Mossburg, Andrew Adezio, Peter Noel, Elizabeth M. Lennon, David P. Cormode

## Abstract

Ulcerative colitis (UC) is a chronic inflammatory bowel disease (IBD). CT imaging with contrast agents is commonly used for visualizing the gastrointestinal (GI) tract in UC patients. CT is a common imaging modality for evaluating IBD, especially in patients with acute abdominal pain presenting to emergency departments. CT’s major limitation lies in its lack of specificity for imaging UC, as the commonly used agents are not well-suited for inflamed areas. Recent studies gastrointestinal tract (GIT) in UC. Further systemic research is needed to explore novel contrast agents that can specifically image disease processes in this disease setting.

## Introduction

Inflammatory bowel disease (IBD) is a heterogenous set of chronic inflammatory between the intestine and the skin, necessitating repeated imaging studies. Delayed intervention can result in fatal consequences, making early detection from contrast-enhanced CT imaging and prompt treatment for IBD and its complications crucial.^3,4^

The diagnosis of IBD involves a comprehensive assessment of clinical findings, inflammatory biomarkers, imaging results, and endoscopic biopsies.^3,5,6^ Due to its swift scan times, ability to provide high-resolution assessments of both intestinal and extra-intestinal disease manifestations, and widespread accessibility in most hospitals around the clock, CT is a common imaging modality for evaluating IBD, especially in patients with acute abdominal pain presenting to emergency departments.^7,8^ In one Canadian study, CT was performed in 17-22% of IBD patients presenting to emergency rooms.^9^ CT is an X-ray based, whole body imaging technique that is relatively inexpensive, simple to use, and fast (image acquisition time <5 s). The speed of CT systems makes imaging of IBD patients easier by minimizing motion artifacts from bowel movement.^7^ Contrast-enhanced CT is useful for investigating extraintestinal manifestations, especially in patients presenting to the emergency department with acute abdominal pain, to determine the presence of potential fistulas.^4,10^ Additionally, CT can be used to analyze patients with Magnetic resonance imaging (MRI)-incompatible devices or metallic foreign bodies.

One major drawback of CT is that the commonly used agents for imaging the gastrointestinal tract in the clinic, such as iodinated small molecules and barium sulfate suspensions, are not ideal for imaging UC due to the lack of specificity for inflamed areas, patient allergic reactions, and the potential for acute kidney injury.^11,12^ Furthermore, there are currently no contrast agents specifically designed for imaging IBD.^13^ Thus, there is a need to develop novel CT contrast agents for improved IBD imaging.^14^

Engineered nanoparticles (NPs) with diverse characteristics hold promise for potential advancements in the diagnosis, treatment, and theranostics of IBD.^15–18^ An example of a novel CT contrast agent for IBD imaging is dextran-coated cerium oxide NPs. These particles were shown to preferentially accumulate at IBD inflammation sites, providing excellent CT contrast in the intestine. The use of dextran in the coating, which has an affinity for inflammatory disease sites, allows for precise localization of colitis.^11^ Despite the potential of NP in advancing the diagnosis, treatment, and theranostics of IBD, a noticeable research gap remains that requires attention. Specifically, the impact of NP size on their behavior and performance in IBD-related applications has not yet undergone systematic study.

Gold nanoparticles (AuNPs) are well known to be effective CT contrast agents, and can be synthesized across a wide range of sizes, offering researchers a versatile toolkit for tailoring properties to specific imaging needs.^19–22^ This aspect makes AuNPs an excellent model system for investigating the impact of NP size on imaging IBD with CT. Therefore, to investigate the impact of AuNP size on CT contrast production, pharmacokinetics, and biodistribution in IBD, we present the properties of five AuNP preparations with sizes ranging from 5 to 75 nm. To explore the impact of AuNP size on CT contrast within a living system, we introduced AuNPs into mice through gavage and conducted imaging at different intervals, alongside biodistribution analysis. Our results indicate that the size of nanoparticles plays a role in diffusion and the creation of contrast within distinct parts of the gastrointestinal tract.

## Methods

### Materials

The following chemicals and materials were acquired from their respective sources. Gold chloride trihydrate (99.9%), hydroquinone (99.9%) sodium citrate dihydrate, and sodium borohydride powder (98.0%) were obtained from Sigma-Aldrich (St. Louis, MI). The clinically-approved iodinated contrast agent, iopamidol (Isovue-300), was acquired from Bracco Diagnostics (Monroe, NJ). 5 kDa m-PEG-thiol was sourced from Creative PEGWorks (Winston Salem, NC, USA). C57BL/6 J mice were procured from The Jackson Laboratory (Bar Harbor, ME).

### Gold Nanoparticle Synthesis

Spherical AuNPs with sizes ranging from 5 to 75 nm were synthesized using the Turkevich method, a modified Turkevich method or a seeded growth method.^23,24^ For the synthesis of 5 nm gold nanoparticles (AuNPs), a solution was prepared using a modified version of the Turkevich method, dissolving 1.6 mL of 1% w/v gold (III) chloride (HAuCl_3_) in 100 mL of ultrapure water. Simultaneously, a 1% w/v solution of sodium borohydride was freshly prepared in cold ultrapure water. The reduction process began by slowly adding 1 mL of the 1% sodium borohydride solution, dropwise, to the HAuCl_3_ solution while continuously stirring. The solution was stirred for 20 minutes, after which ligand exchange was performed (see procedure noted below).^24^

AuNPs with diameters of 15 nm and 28 nm were produced using the Turkevich method. In brief, a solution consisting of 600 μL of 1% w/v HAuCl_3_ in 60 mL of deionized water was heated to boiling, after which 1.8 mL of 1% w/v sodium citrate dihydrate was added. The solution was then continuously heated and stirred for 10 minutes before being allowed to cool to room temperature. Subsequently, the solution was allowed to stir at room temperature for three hours before proceeding with the ligand exchange process.^23^

AuNPs with diameters of 51 nm and 75 nm were produced using a seeded growth method. 15 nm AuNP seeds were prepared using the above described method. 1 mL of 1% w/v HAuCl_3_ was mixed with 100 mL of deionized water at room temperature while vigorously stirring. Subsequently, an appropriate volume of 15 nm seeds was added to the solution to achieve the desired nanoparticle size (3 mL for 51 nm and 750 µL for 75 nm). Following the addition of the seeds, 220 μL of 1% w/v sodium citrate dihydrate was introduced to the mixture, followed immediately by 1 mL of 0.03 M hydroquinone. The solution was left to stir for three hours at room temperature prior to the ligand exchange process.^25^

### Ligand Exchange and Purification

Ligand exchange was carried out using 5 kDa thiolated polyethylene glycol (PEG-SH). PEG was first dissolved in water to achieve a concentration of 5 mg/mL. The amount of PEG-SH differed depending on the size of the AuNP, as per previous reports.^23,25^ The PEG ligands were then added in varying amounts depending on the size of the gold nanoparticles. Specifically, we added 2.522 mL for 5 nm, 341.41 µL for 15 nm, 204.84 µL for 28 nm, 177.05 µL for 51 nm, and 114.86 µL for 75 nm.^23,25^ Subsequently, the solution was stirred for 6 hours at room temperature.

Following ligand exchange, all AuNPs underwent purification and concentration using centrifugation. For 5, 15, and 28 nm AuNPs, the particles were initially purified three times using 10 kDa molecular weight cutoff (MWCO) tubes with deionized water. Each purification step involved centrifugation at a speed of 2500 rcf for 10 minutes. Subsequently, 10 mL of phosphate buffered saline (PBS) was added to the same tube, and the AuNPs were concentrated to 1 mL by centrifuging at the same speed for an additional 15 minutes. The concentrated AuNPs were then collected. 51 and 75 nm AuNPs were centrifuged at a speed of 2000 rcf for 20 minutes in a 50 mL falcon tube. The supernatant was discarded, and deionized water was added to restore the solution’s volume to 50 mL. This purification step was repeated twice, followed by collection of the AuNPs, which were transferred to 1.5 mL Eppendorf tubes. PBS was added to bring the total volume to 1.5 mL, and the tubes were centrifuged at the same speed as before. After centrifugation, the supernant was discarded, leaving behind the concentrated AuNP product. For all sizes, the concentrated AuNPs were collected and passed through a 0.45 μm syringe filter to a 1.5 mL eppandord tube.

### Nanoparticle Characterization

The AuNP formulations were characterized using transmission electron microscopy (TEM) (JEOL 1010 electron microscope), zeta potential, and dynamic light scattering (DLS) (Zetasizer Nano ZS90). The Au concentrations were determined through inductively coupled plasma-mass spectrometry system (ICP-OES) (Spectro Genesis ICP, Kleve, Germany). Each nanoparticle batch was characterized for size with TEM and ultraviolet–visible (UV-vis) spectroscopy to ensure the consistency of the formulations.

### Transmission Electron Microscopy

To analyze the size and shape of the AuNP cores, transmission electron microscopy (TEM) was utilized. A JEOL 1010 microscope from JEOL USA Inc. (Peabody, MA) operating at 80 kV, was employed for this purpose. 10 µL of diluted AuNP solution was deposited on carbon-coated copper grids (FCF-200-Cu, Electron Miccroscopy Services, Hatfield, PA) and allowed to evaporate before imaging. The size of the AuNP cores was determined by manually measuring the particles using imageJ software developed by the National Institutes of Health.

### UV/visible Absorption

UV-vis spectra were recorded using an Evolution 201 UV-visible spectrophotometer from Thermo Scientific (Waltham, MA), which had a wavelength range of 200 nm to 800 nm. To perform the measurement, 2 μL of the AuNP stock solution was mixed with 998 μL of deionized water. For each measurement, a cuvette was filled with 1 mL of the diluted AuNP formulation, and absorbance was measured.

### Dynamic Light Scattering and Zeta Potential

The hydrodynamic diameters and zeta potential of the AuNPs were determined using a Nano ZA-90 Zetasizer from Malvern Instruments (Worcestershire, UK). To perform the measurements, 995 μL of deionized water was added to 5 μL of the AuNP stock solution for dilution. Subsequently, 1 mL of the diluted solution was transferred into a cuvette for analysis. The hydrodynamic diameter was reported as the Z-average, which represents the mean diameter of the particles in the solution accounting for their Brownian motion.

### Inductively Coupled Plasma Optical Emission Spectroscopy

Concentrations of the AuNP solutions were determined using ICP-OES with a Genesis ICP instrument from Spectro Analytical Instruments GmbH (Kleve, Germany). To prepare the samples, four different samples of 5 μL of the AuNP stock solution were dissolved in 995 μL of aqua regia. After 2 hours, 5 mL of deionized water was added to each solution, bringing the final volume to 6 mL. The concentrations of gold in these samples were then measured. The concentration of AuNP in the stock solution was calculated by averaging the values obtained from the four samples, adjusting for dilution.

### *In Vitro* Phantom Imaging

The X-ray contrast generation capabilities of the AuNP formulations were assessed using a conventional clinical CT system. The samples were scanned in custom-made plastic phantoms to imitate the beam hardening effect in patients.^23^ Each AuNP size, represented at a range of concentrations, was meticulously placed in vials and arranged in a plastic rack. To replicate the conditions of a human abdomen, the rack with vials was immersed in a plastic container filled with water to a height of 21 cm before conducting clinical CT phantom imaging. This setup allowed for a realistic representation of the CT scanning environment and provided valuable data on the contrast properties of the AuNPs in simulated clinical conditions.^23^ The samples were prepared in triplicates and at concentrations of 0.0, 0.5, 1, 2, 4, 8, and 12 mg/mL. The contrast produced from these formulations was compared with that of iopamidol and water. The conventional CT contrast properties of AuNP were evaluated by scanning samples using a Spectral CT 7500, Philips Healthcare, Best, Netherlands scanner at 80, 100, 120, and 140 kV. In each case, a matrix size of 512 x 512, a field of view of 37 x 37 cm, and a reconstructed slice thickness of 0.5 mm was used. T-values were calculated from the attenuation rates and standard errors of the data. These values were compared to the reference t-value of 2.45, using six degrees of freedom and a 5% significance level.

### Animal Models

These procedures were approved by the Institutional Animal Care and Use Committee (IACUC) of the University of Pennsylvania under protocol number 806566. Acute colitis was induced in mice using dextran sodium sulfate (DSS) following a previously published protocol.^26,27^ On Day 1, C57BL/6 mice, aged 7 weeks, were weighed and assigned identification numbers. The water supply in the mouse cages was filled with a 3% (w/v) solution of DSS obtained from MP Biomedicals (Irvine, CA). This DSS solution, with a molecular weight range of 36,000 to 50,000 Da, was used to induce IBD. On days 3 and 5, the remaining DSS solution in the water bottles was removed, and fresh DSS solution was added. Finally, on Day 8, the remaining DSS solution was replaced with autoclaved water to discontinue DSS exposure.^26,27^ Age-matched, untreated mice were used as healthy controls. The study considered sex as a biological variable and used mice in a 1:1 male-to-female ratio, as IBD affects both sexes.^28^

### *In Vivo* CT Imaging

The previously described mice underwent CT scans before and immediately following the administration of the agents (5 min.) to assess the generation of CT contrast. Additional scans were performed at 1, 2, and 24 hours post-administration. The shorter time points were selected as they are clinically relevant, while the longer time point was chosen since it is expected that most of the nanoparticles would be excreted from the body within 24 hours. The sample size was 6 per group. The dose for AuNP and iopamidol was 250 mg Au or I/kg body weight (typical doses for x-ray contrast agents). All treatments were volume-matched and given via gavage. The mice were scanned with a MILabs U-CT (MILabs, Utrecht, The Netherlands)scanner using a tube voltage of 55 kV and isotropic 100 micron voxels. The images were analyzed using Osirix MD software. The change in attenuation values between pre- and post-administration time points was measured in the gastrointestinal tract and organs of interest such as the liver, kidney, and spleen. To assess whether there were significant differences in attenuation between different AuNP formulations and the control groups, a Tukey’s multiple comparison test with a single pooled variance was employed. This statistical test was utilized to compare the means of multiple groups and determine if there were significant differences among them.

### Biodistribution

48 female and 48 male mice were euthanized using anesthesia and cervical dislocation. Organs including the stomach, small intestine, large intestine, as well as vital organs such as the heart, liver, and spleen, along with blood, were collected and weighed. These samples were digested in 6 mL of nitric acid (for liver, spleen, and kidneys) and incubated overnight at 90°C. The following day, the samples were diluted to a final volume of 10 mL with DI water. Gold nanoparticle concentrations were quantified using ICP-OES, and biodistribution data is reported as mean %ID/g ± standard error of the mean (SEM).

### Statistical Methods

The figures in the study include error bars representing the standard error from the mean, while Table 1 presents mean and standard deviation of AuNP core diameter and zeta potential for different AuNP sizes. For examining the interactions between each AuNP size in the biodistribution of gold content in organs, a Tukey’s multiple comparisons test was conducted. This statistical test enabled the comparison of different AuNP sizes and identified any significant differences in the distribution of gold content among organs. All statistical analyses were performed using Graphpad Prism 7 software (GraphPad Prism Software Inc., San Diego, California).

**Table 1.** Summary of the physical properties of the AuNP used in this study. Core sizes were determined using TEM, while hydrodynamic diameters and zeta potential were measured using DLS. Absorbance values were obtained using a UV-vis spectrometer. The plus-minus symbol indicates the standard deviation of the measurements for each parameter.

| Core diameter (nm) | Hydrodynamic diameter (nm) | Zeta-potential (mV) | Absorbance peak (nm) |
| --- | --- | --- | --- |
| 5.3 ± 1.1 | 20.0 ± 0.3 | -9.0 ± 0.7 | 516 |
| 14.5 ± 1.9 | 41.1 ± 0.8 | -20.5 ± 1.0 | 524 |
| 28.4 ± 4.3 | 46.0 ± 0.8 | -22.0 ± 0.7 | 528 |
| 50.6 ± 5.1 | 97.5 ± 0.5 | -26.4 ± 1.1 | 536 |
| 75.2 ± 7.0 | 106.7 ± 2.3 | -29.0 ± 0.7 | 552 |

## Results

### Synthesis and Characterization

To evaluate their properties as CT contrast agents and for imaging the GI tract, AuNPs were synthesized with core diameters ranging from 5 to 75 nm. In order to improve their stability, the AuNPs were subsequently coated with PEG, a widely used capping ligand that results in excellent stability and biocompatibility in both our laboratory and those of others.

TEM was utilized to examine the size and shape of the AuNP cores, as shown in **Fig. 1**. The majority of the observed AuNPs exhibited a spherical shape, and there was minimal variation in size for a given sample, indicating low polydispersity among the particles. Table 1 summarizes the core size measurements obtained from the micrographs, indicating successful control over AuNP size using our synthetic methods. ^25,29^ In this manuscript, the formulations are referred to based on their core diameters. This allows for clear identification and differentiation of the different formulations throughout the document.

**Figure 1.**
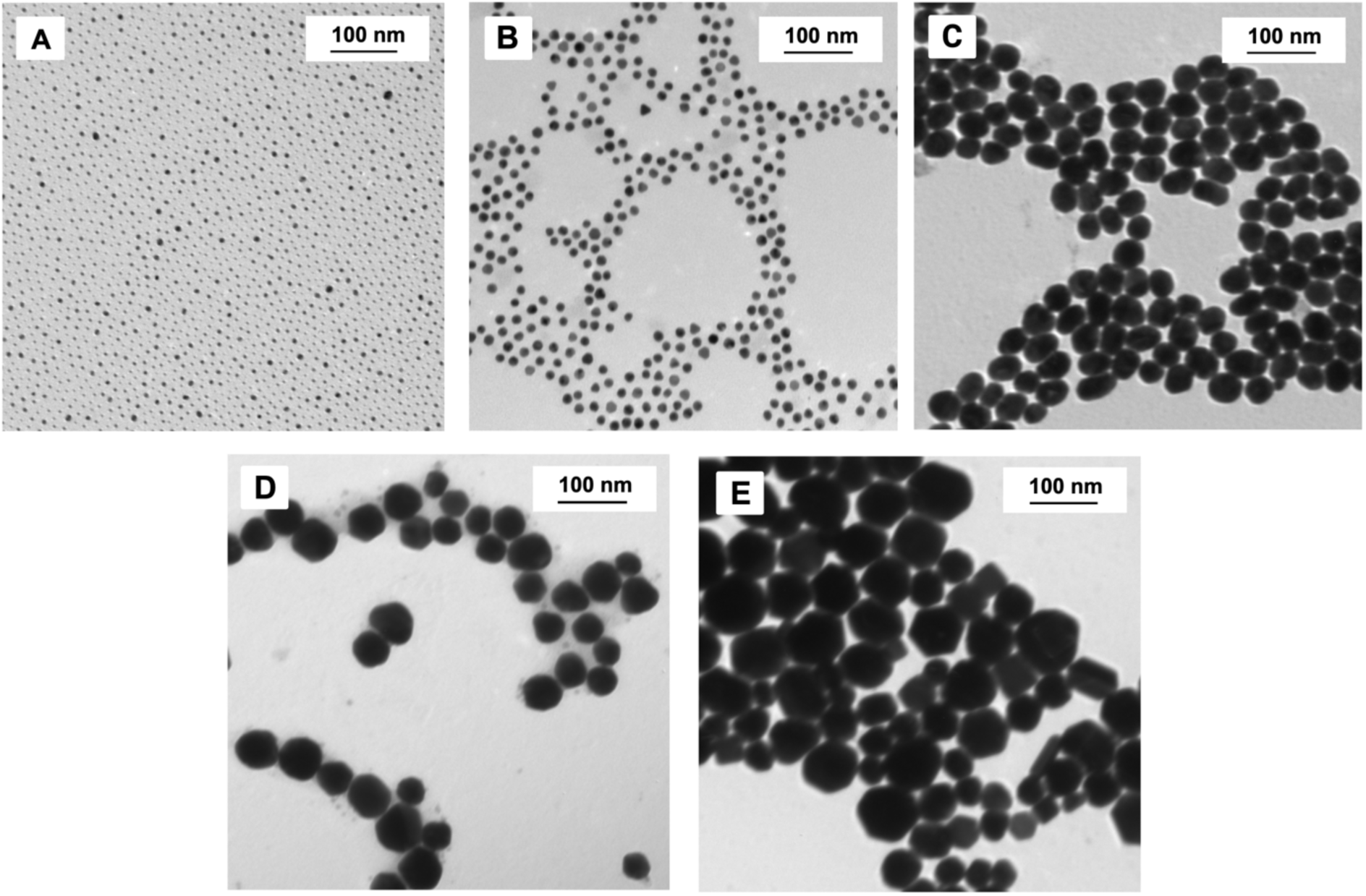
Representative TEM micrographs of the different AuNP formulations, with each panel corresponding to a specific size: **(A)** 5, **(B)** 15, **(C)** 28, **(D)** 51, and **(E)** 75 nm. In all panels, the scale bars indicate a length of 100 nm.

The absorption peaks of the AuNPs occurred from 516 to 552 nm, with increases in the wavelengths of the peaks as the diameters of the gold nanoparticles increased, as expected.^30^ The hydrodynamic diameters of the AuNPs were measured and found to be slightly larger than that of the core diameter, reflective of PEG coating. (**Table 1**).

### CT Contrast Properties

We and others have previously reported no effect of AuNP size on CT contrast generation.^23^ To ensure accurate interpretation of our *in vivo* experiments, we investigated the CT contrast generating properties of the AuNP reported herein via CT phantom imaging. A phantom containing six different concentrations of AuNPs for each size was prepared in triplicate. To investigate the CT contrast properties, the phantom was scanned using a clinical scanner at four different tube voltages (80, 100, 120, and 140 kV). To simulate the beam hardening experienced by patients, we surrounded the samples with 21 cm of water. Representative CT phantom images of an AuNP formulation are displayed in **Fig. 2A** (specifically, 15 nm using 80 kV).

**Figure 2.**
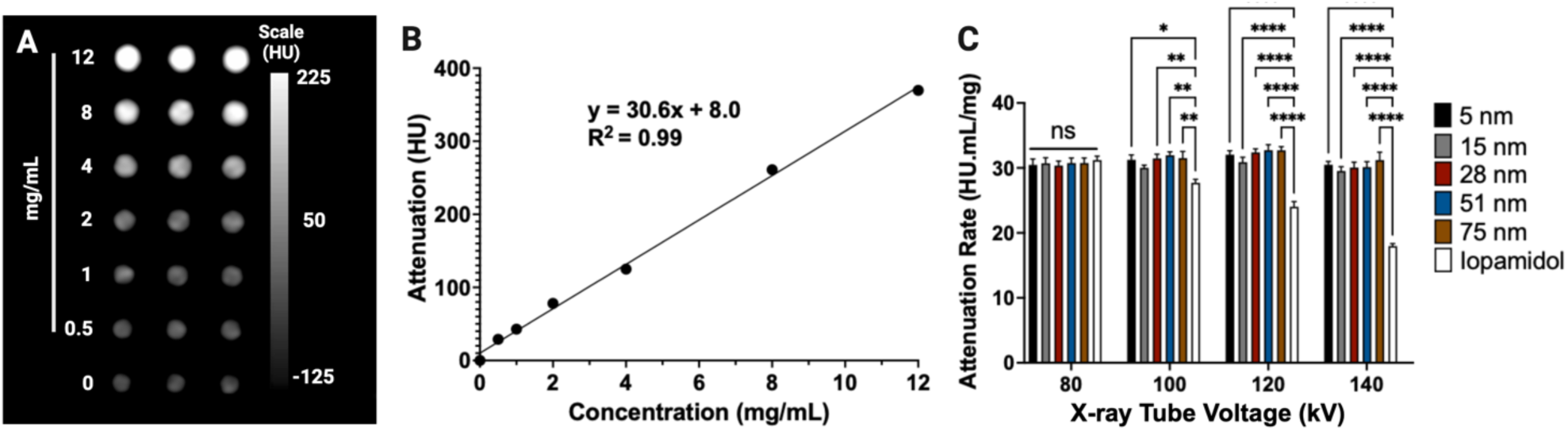
(A) Representative CT images of a phantom containing 15 nm AuNP. **(B)** X-ray attenuation as a function of concentration for 28 nm AuNP scanned by CT at 80 kV. **(C)** Attenuation rate values for AuNP formulations derived from CT scans performed at 80, 100, 120, and 140 kV. Error bars represent standard error of the mean, * indicates P < 0.05, ** indicates P < 0.01, and **** indicates P < 0.0001.

As expected, we found that there were linear correlations (R^2^ = 0.99) between the CT attenuation of AuNP and the mass concentration of Au for each tube voltage used (**Fig. 2B**). Importantly, our findings demonstrated no statistically significant differences in x-ray attenuation between the various AuNP sizes when using clinical CT systems (**Fig. 2C**). These results are consistent with our previous studies, reinforcing the notion that the size of AuNPs does not have a significant impact on x-ray attenuation.^23,31^ This has the benefit that any differences observed in attenuation *in vivo* from the different AuNP will arise solely from differences in agent location as opposed to contrast generation.

When compared with iopamidol, a commercially available contrast agent, we found that AuNP and iopamidol had similar attenuation rates at 80 kV, but AuNP produced a slightly greater CT attenuation rate at 100 kV. However, AuNP had a significantly higher attenuation rate than iopamidol at 120 and 140 kV, as gold has a higher k-edge value than iodine. These results are also consistent with prior findings from our group and others.^31–33^

### *In Vivo* CT Imaging

The utility of the above described AuNPs for GI tract imaging was assessed through the use of a MILabs U-CT system. In the study, both healthy mice and mice with DSS-induced colitis were used, and AuNPs or iopamidol were administered. The mice underwent pre-scans, followed by the administration of AuNPs of different sizes via gavage at a concentration of 30 mg/mL. The mice were then imaged at different time points. In line with our expectation and prior reports, strong CT contrast was observed in the GI tract of these mice, with no contrast apparent in other organs.^11,34^ **Fig. 3** and **Fig. S1-S3** present CT images of colitis mice before and after administration with different sizes of AuNPs, demonstrating strong contrast in various sections of the gastrointestinal tract (GIT) at different time intervals.

**Figure 3.**
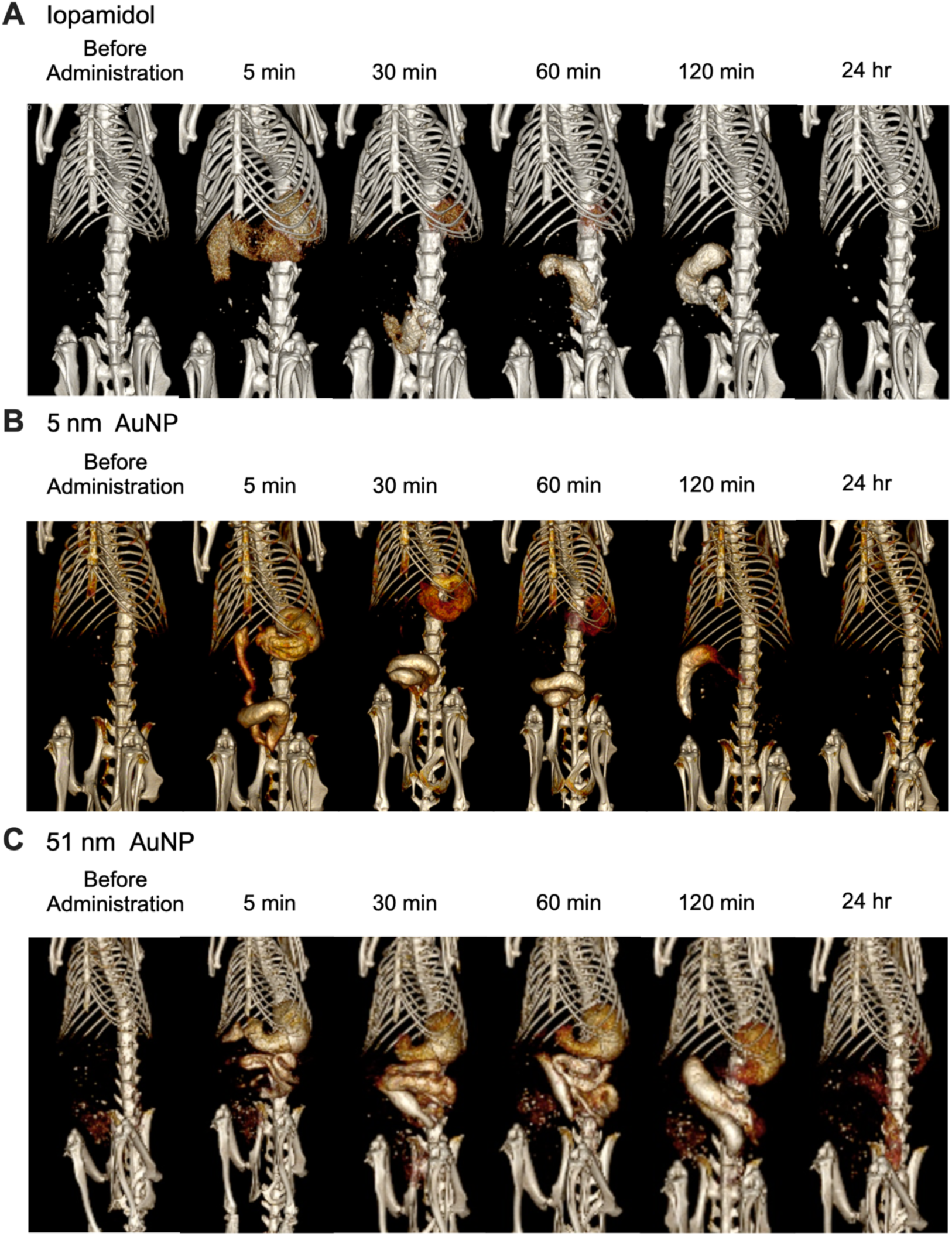
Representative CT images of mice with colitis before and at the timepoints noted after oral administration of AuNPs: **(A)** Iopamidol **(B)** 5 nm **(C)** 51 nm. Images are displayed at a window level of 1090 HU and window width of 930 HU.

*In vivo* imaging studies demonstrated that the attenuation of the nanoparticle formulations in the control mice decreased with the passage of time for all particle sizes in the stomach (**Fig. 4A, 4D**), which is as expected since the agents are deposited there with minimal absorption and then disperse further into the GI tract. In the small intestine, the attenuation was highest at 60 minutes (**Fig. 4B, 4E**). On the other hand, the highest attenuation in the large intestine for each tested agent was observed at the 120 minute time point (**Fig. 4C, 4F**). Interestingly, our results revealed that the attenuation of iopamidol was lower than that of the AuNP formulations, which is likely due to lower inherent contrast generation from iodine (**Fig. 2**) and possibly diffusion of the agent out of the intestine into other tissues. The results of CT imaging in mice with colitis follows a similar trend to the control group, consistently revealing a decrease in attenuation in the stomach as time progressed for all particle sizes, highest attenuation at the 60 minutes in the small intestine and, in the large intestine, a continuous increase in attenuation from 0 to 120 minutes.

**Figure 4.**
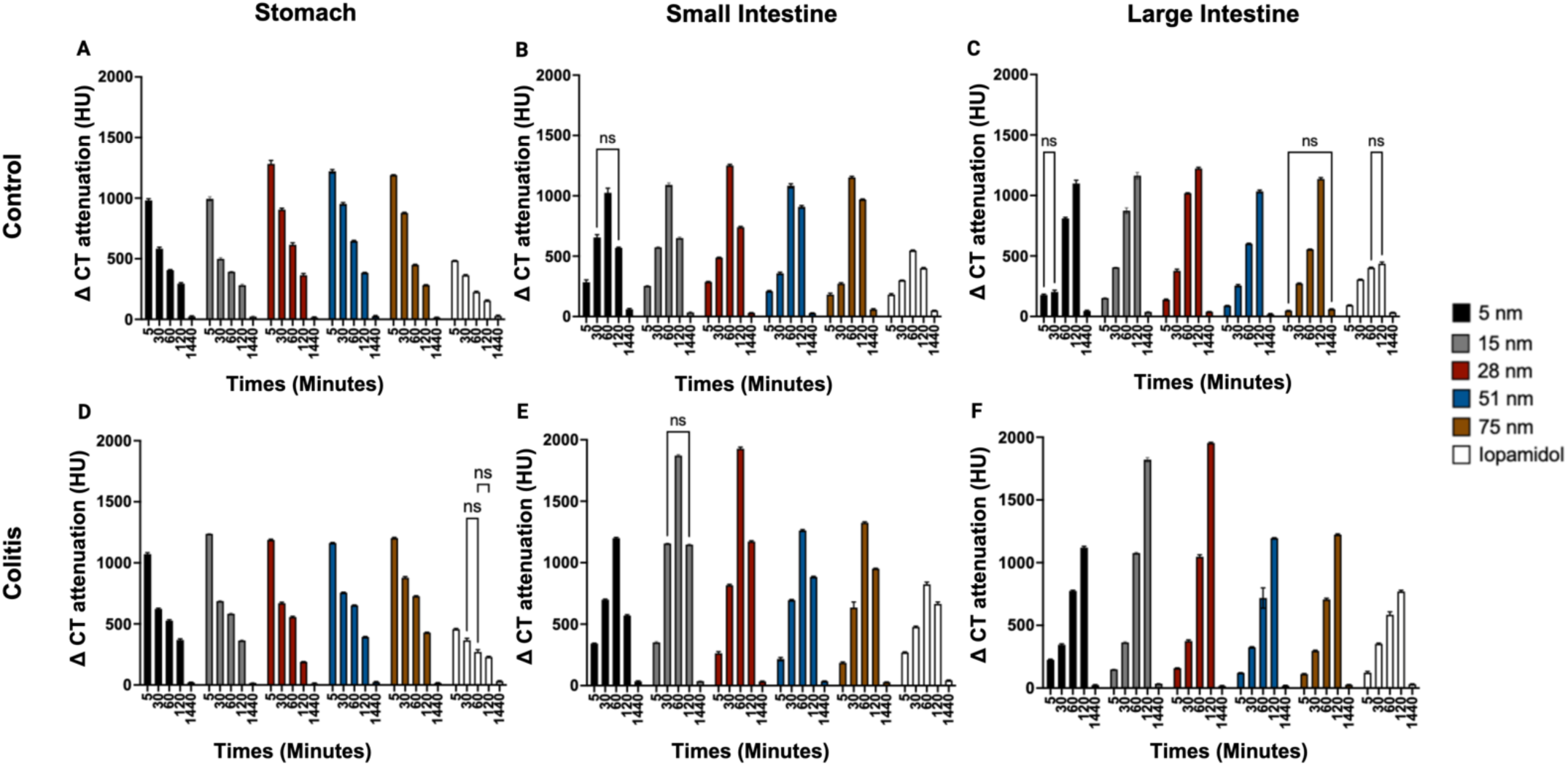
Attenuation dynamics of AuNP (5-75 nm) and iopamidol in the GI tract of different organs over time, including the stomach (**A**), small intestine (**B**) and large intestine (**C**) of control mice, and stomach (**D**), small intestine (**E**) and large intestine (**F**) of colitis mice. Error bars represent standard error of the mean. Only non-significant p values are displayed on the graph for clarity. All P values, including significant ones, can be found in the Supplemental Table S1-S6.

The results from both the control and colitis mice demonstrate some differences in attenuation and contrast among the nanoparticle formulations, as well as iopamidol, in different sections of the gastrointestinal tract at specific time points (**Fig. 3, 4, S1-S4**). For example, in the control mice, the 28 nm nanoparticles exhibited somewhat greater attenuation than the other nanoparticles at the 5-minute timepoint (**Fig. 4, S4**). In addition, in the small intestine, the 28 nm AuNP formulation produced more attenuation than the other formulations at the 60 minute timepoint (**Fig. 4, S4**). However, overall there were no dramatic differences in the contrast produced between the different AuNP sizes tested.

When comparing the control group with the colitis group, differences in agent behavior were observed. For example, the agents seemed to clear more slowly from the stomach and higher attenuation values were noted in small intestine, and large intestine, perhaps as a result of the impeded gastric function found in this disease.

### GI tract retention

We conducted a study to examine the retention of AuNP in healthy and DSS-induced colitis mice 24 hours after oral administration. The concentrations of gold in the stomach, small intestine, large intestine, spleen, heart, and liver were determined using ICP-OES and are shown in **Fig. 5** and **S8.**

**Figure 5.**
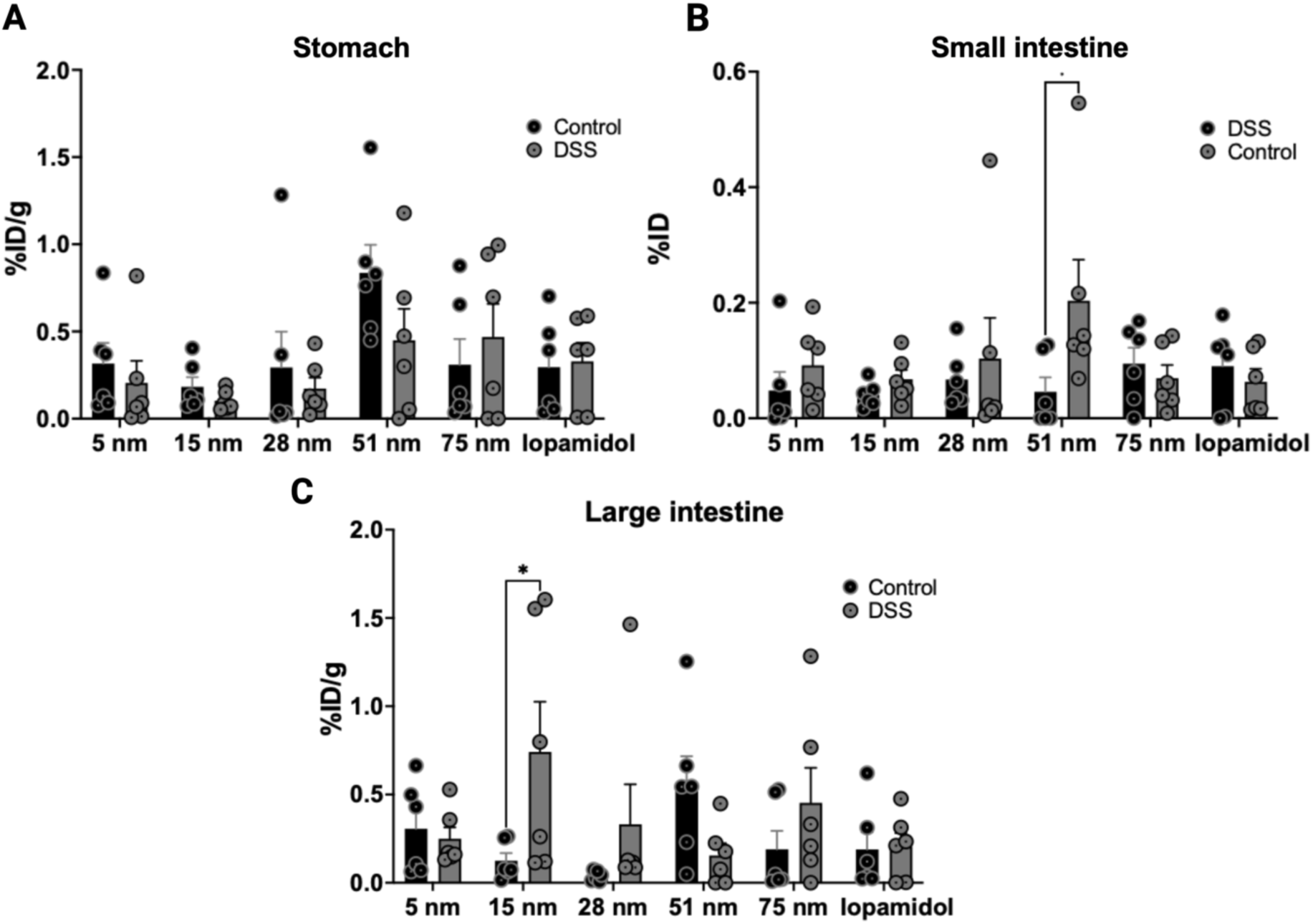
Retention of AuNPs in the organs of the GI tract in healthy and DSS-colitis mice 24 hours after administration. %ID/g values in (**A**) the stomach, (**B**) small intestine, (**C**) and large intestine. Error bars represent standard error of the mean, and * indicates P < 0.05.

Our findings indicate that the biodistribution of AuNP was similar in healthy and colitis mice. In healthy mice, the quantity of gold found in the stomach, small intestine, and large intestine was relatively low and less than 0.7% ID/g was retained, indicating nearly complete clearance of AuNP at 24 hours, consistent with *in vivo* CT imaging results (**Fig. 4**). In colitis-induced mice, small amounts of gold were also detected in the stomach, small intestine, and large intestine (less than 0.6% ID/g) (**Fig. 5).** Furthermore, a negligible amount of gold was found in the heart, liver, and spleen of both colitis and control animals (less than 0.8% ID/g).

## Discussion

In this study, we successfully synthesized PEGylated AuNP with gold core diameters between 5 and 75 nm. Importantly, our experiments demonstrated that the size of AuNPs does not affect their inherent CT contrast properties. This suggests that AuNPs of different sizes can be used interchangeably as contrast agents for CT imaging, when contrast production alone is considered. Our experiments also compared the CT attenuation rates of AuNPs and iopamidol, a commonly used contrast agent, which showed that AuNPs had comparable or slightly greater attenuation rates than iopamidol at lower tube voltages (80 and 100 kV), but had significantly higher attenuation rates than iopamidol at higher tube voltages (120 and 140 kV) due to the higher k-edge value of gold compared to iodine.^35^ These findings were consistent with prior results.^23,36,37^ *In vivo* imaging using a MILabs µ-CT system was used to evaluate the pharmacokinetics and biodistribution of orally administered AuNPs of different sizes in healthy mice and mice with DSS-induced colitis. Our findings indicate a general decrease in attenuation within the stomach over time, observed in both the control and colitis groups. Conversely, in the small intestine and large intestine, attenuation increased progressively until reaching its peak, occurring at 60 and 120 minutes, respectively.

We found variations in attenuation and contrast across the AuNP formulations and iopamidol within segments of the gastrointestinal tract at specific time intervals. For the control animals, the 28 nm formulation produced slightly greater attenuation in each organ at certain time points.

Notably, iopamidol yields lower contrast compared to other formulations due to differences in contrast generation and potential diffusion from the gastrointestinal tract into the body of the mouse.^11^

We typically observed greater maxiumum attenuation values in the colitis group compared with the control group. This is perhaps due to nanoparticle interactions with the mucus layer lining the GIT facing altered dynamics in the presence of colitis—a condition characterized by inflammation of the colon. The inflamed environment associated with colitis disrupts the physiological conditions and barrier functions of the GIT.^17^ This disruption has multifaceted effects on nanoparticle behavior, including heightened permeability of the intestinal barrier, potentially facilitating easier absorption into inflamed tissues.^38^ Additionally, the compromised mucus barrier in colitis may lead to varied nanoparticle interactions, potentially enhancing penetration or altering retention times.^39^ Since the AuNP size did not result in substantial effects on imaging results, considerations for which type of nanoparticle to develop can switch to other factors such as cost or ease of synthesis (which, in our opinion, would favor AuNP synthesized by the Turkevich method).

Our previous research involving dextran coated cerium oxide nanoparticles (Dex-CeNP) revealed a remarkable capacity to enhance CT contrast in GIT and to effectively accumulate in tissues affected by colitis.^11^ Based on our prior outcomes with Dex-CeNP, we hypothesized that a similar accumulation pattern might be observed with AuNP in colitis mice, in comparison to the control group.^11^ However, this did not occur. The most plausible explanation for this discrepancy lies in the distinct coatings employed for the nanoparticles. In our earlier investigations, we utilized dextran as the coating material, a choice driven by its potential to effectively target inflammation,^11^ whereas herein we used PEG. It may be the case that dextran coated AuNP would yield results akin to our previous findings, ultimately leading to comparable outcomes.

Importantly, our study showed that, from the perspective of agent retention, the practicality of use of AuNP as a CT contrast agent for GI tract imaging in patients with colitis is not heavily affected by the underlying disease, as there was no significant difference in the biodistribution between healthy and colitis mice. The low levels of gold found in the stomach, small intestine, and large intestine in both healthy and colitis mice indicate that the majority of AuNP were cleared from the body within 24 hours, and the negligible amount of gold found in the heart, liver, and spleen of both healthy and colitis mice suggests that there is minimal systemic distribution of AuNP.This underscores the potential applicability of -AuNP in CT imaging for colitis patients.

However, it is important to acknowledge certain limitations in this study. Firstly, the use of a small animal model of colitis may not fully replicate the complexity and heterogeneity of human disease. While it provides valuable insights, the findings should be interpreted with caution when extrapolating to human patients.^45^ Secondly, although we observed the effects of the tested AuNP sizes on contrast properties and biodistribution, it is possible that sizes outside this range may behave differently. In addition, in this application, we used high doses of inorganic materials. Studies involving lower doses of agents or the use of soft materials may result in different findings. Furthermore, challenges in precisely identifying the small intestine versus the large intestine and potential variability in the severity of colitis among individual mice introduced additional complexities to the analysis.

## Conclusion

In this study, we synthesized and characterized spherical PEG-coated AuNPs with core sizes ranging from 5 to 75 nm. Consistent with prior work, we found no effect of AuNP size on CT contrast generation and that AuNP generated superior contrast to iodinated agents at higher x-ray tube voltages. To investigate the effect of AuNP size on *in vivo* CT contrast, we administered AuNPs via gavage to mice with and without colitis and imaged them at various time points, while also analyzing biodistribution. Surprisingly, our findings suggest that there is little effect of nanoparticle size on diffusion and contrast generation in different regions of the gastrointestinal tract. We did find that AuNP uniformly provided stronger contrast than iopamidol, a commonly clinically used contrast agent. We note that we only tested one nanoparticle coating and type herein, therefore effects of nanoparticle size of that might be found with nanoparticle compositions is uncertain. However, for similar agents, our findings indicate that considerations such as cost or ease of synthesis would be more important for further development. In addition, the negligible retention of the AuNP administered via this route indicates potential for clinical translation. These findings therefore provide valuable insights for the development of contrast agents for gastrointestinal imaging.

## Acknowledgements

We gratefully acknowledge the support of the NIH under grants F31-EB034165 (DNRB), R21-EB029158 (DPC) and R25 CA140116/CA/NCI (SUPERS program). We would like to express our sincere gratitude to Biao Zuo and the Electron Microscopy Resource Laboratory at the University of Pennsylvania for their invaluable assistance with TEM. We are also grateful to Eric Blankemeyer and David Burney for their expert help with micro-CT imaging and ICP-OES, respectively.

