## Supplemental Data 1 for "The effect of the size of gold nanoparticle contrast agents on CT imaging of the gastrointestinal tract and inflammatory bowel disease"

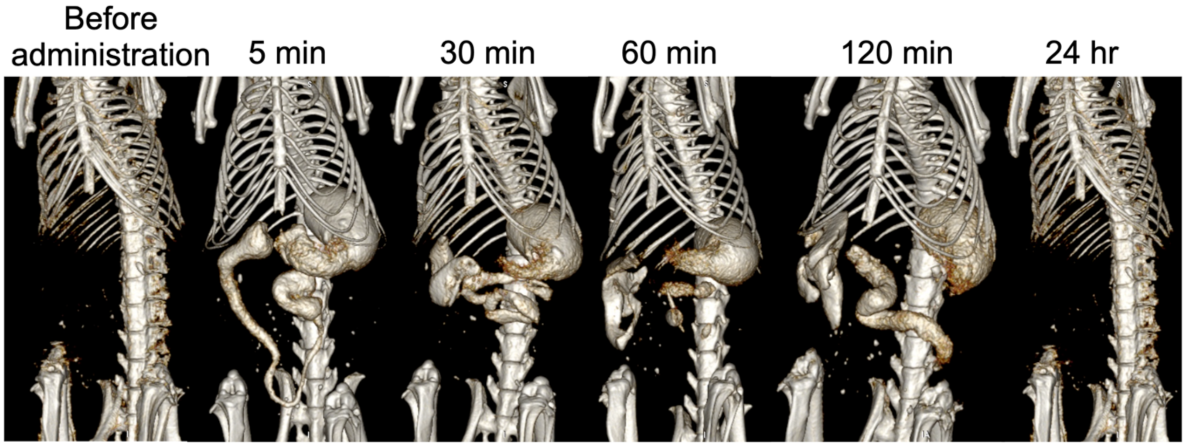


Figure S1: Representative CT images of colitis mice, pre and 24 hrs post oral administration of 15 nm AuNP. Images are displayed at a window level of 1090 HU and window width of 930 HU.

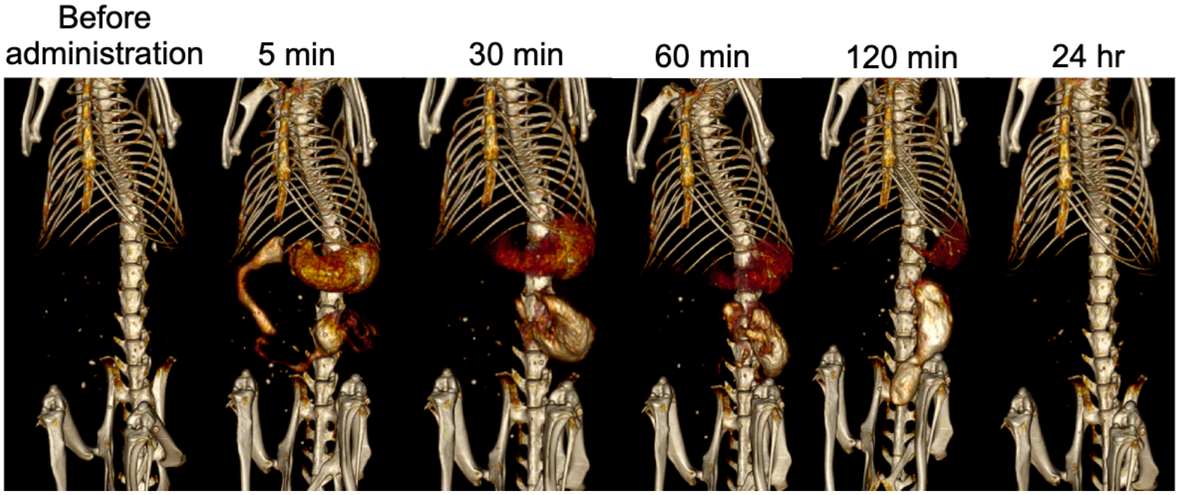


Figure S2: Representative CT images of colitis mice, pre and 24 hrs post oral administration of 28 nm AuNP. Images are displayed at a window level of 1090 HU and window width of 930 HU.

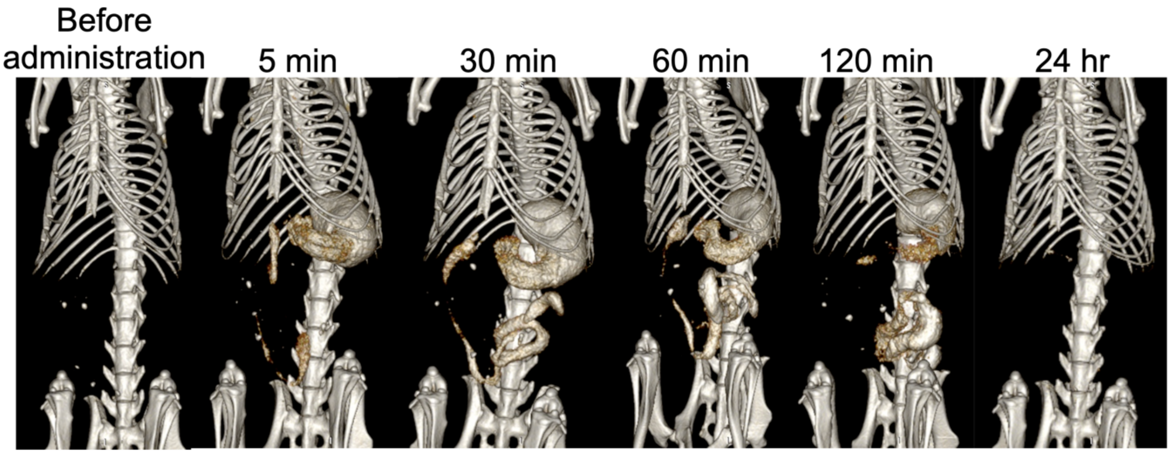


Figure S3: Representative CT images of colitis mice, pre and 24 hrs post oral administration of 75 nm AuNP. Images are displayed at a window level of 1090 HU and window width of 930 HU.

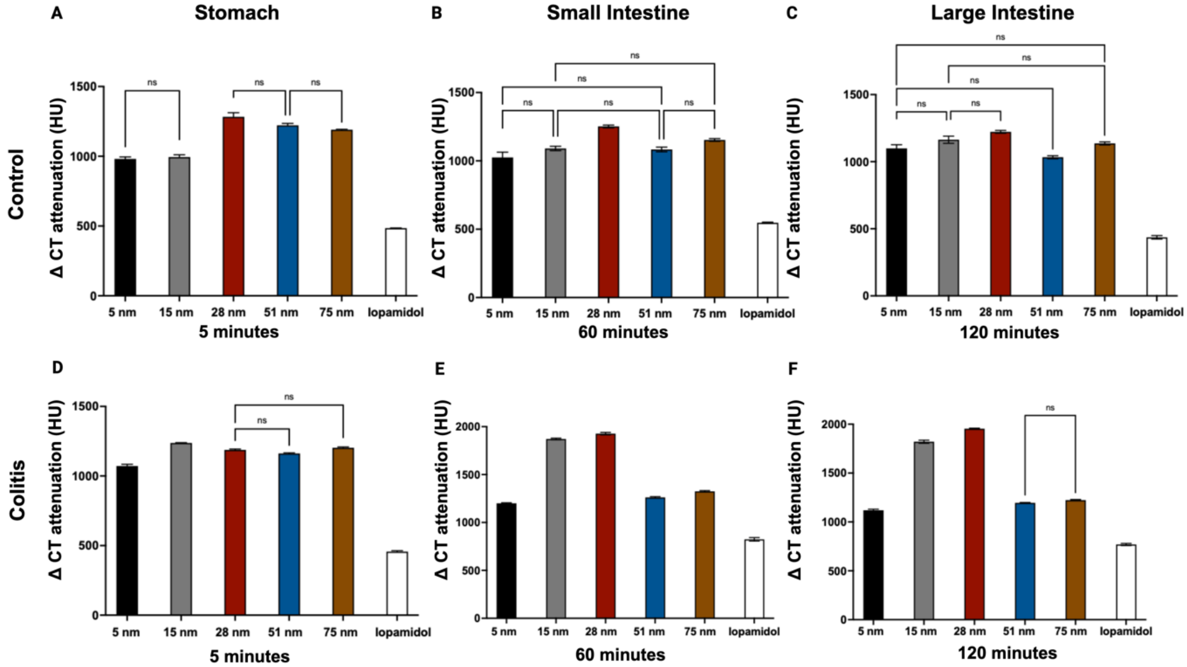


Figure S4: Attenuation dynamics of AuNP (5-75 nm) and iopamidol in the GI tract of different organs. Attenuation values in the stomach at 5 min. (**A**), small intestine at 60 min.(**B**) and large intestine at 120 min. (**C**) of control mice. Attenuation values in the stomach at 5 min. (**D**), small intestine at 60 min. (**E**) and large intestine at 120 min. (**F**) of colitis mice. Error bars represent standard error of the mean, only non-significant P values are displayed on the graph for clarity. All P values, including significant ones, can be found in the Supplemental Table S7-S8.

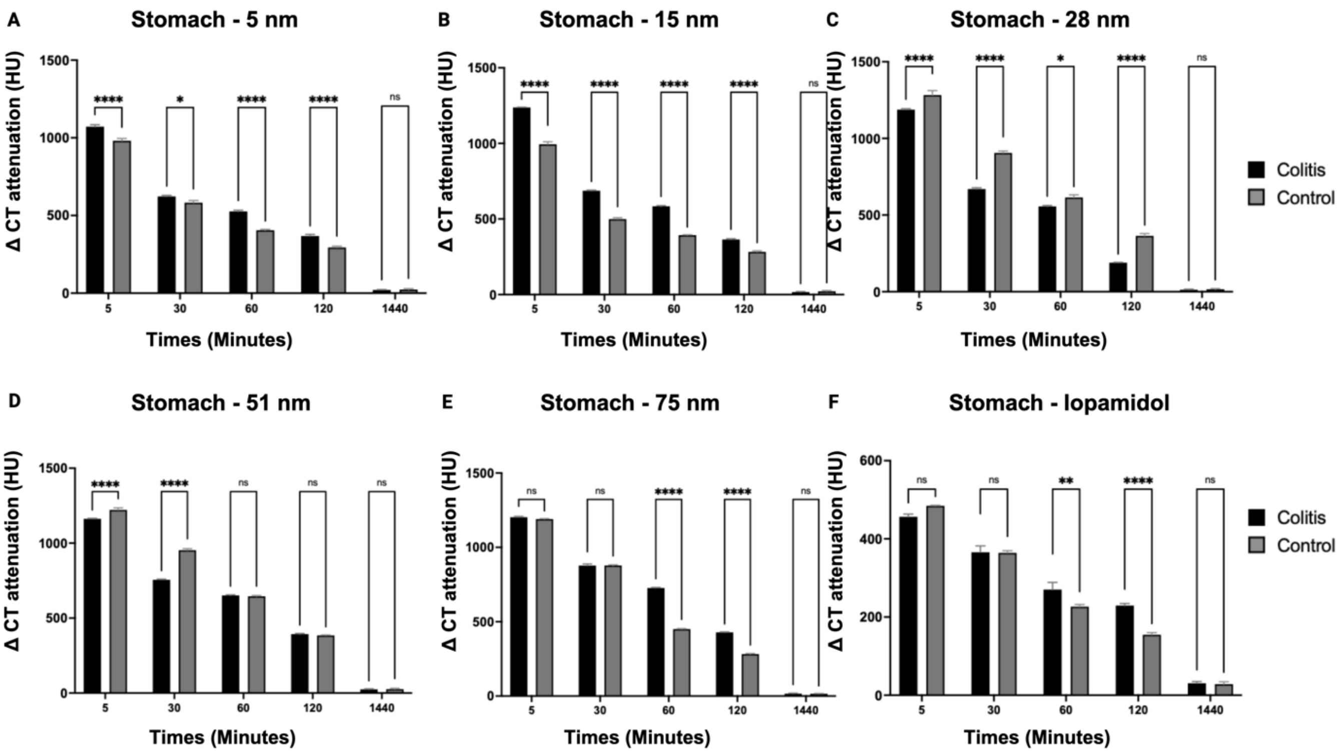


Figure S5: CT attenuation in the stomach from control and colitis mice at different time points. Error bars represent standard error of the mean, * Indicates P < 0.05, ** Indicates P < 0.01, and **** Indicates P ≤ 0.0001.

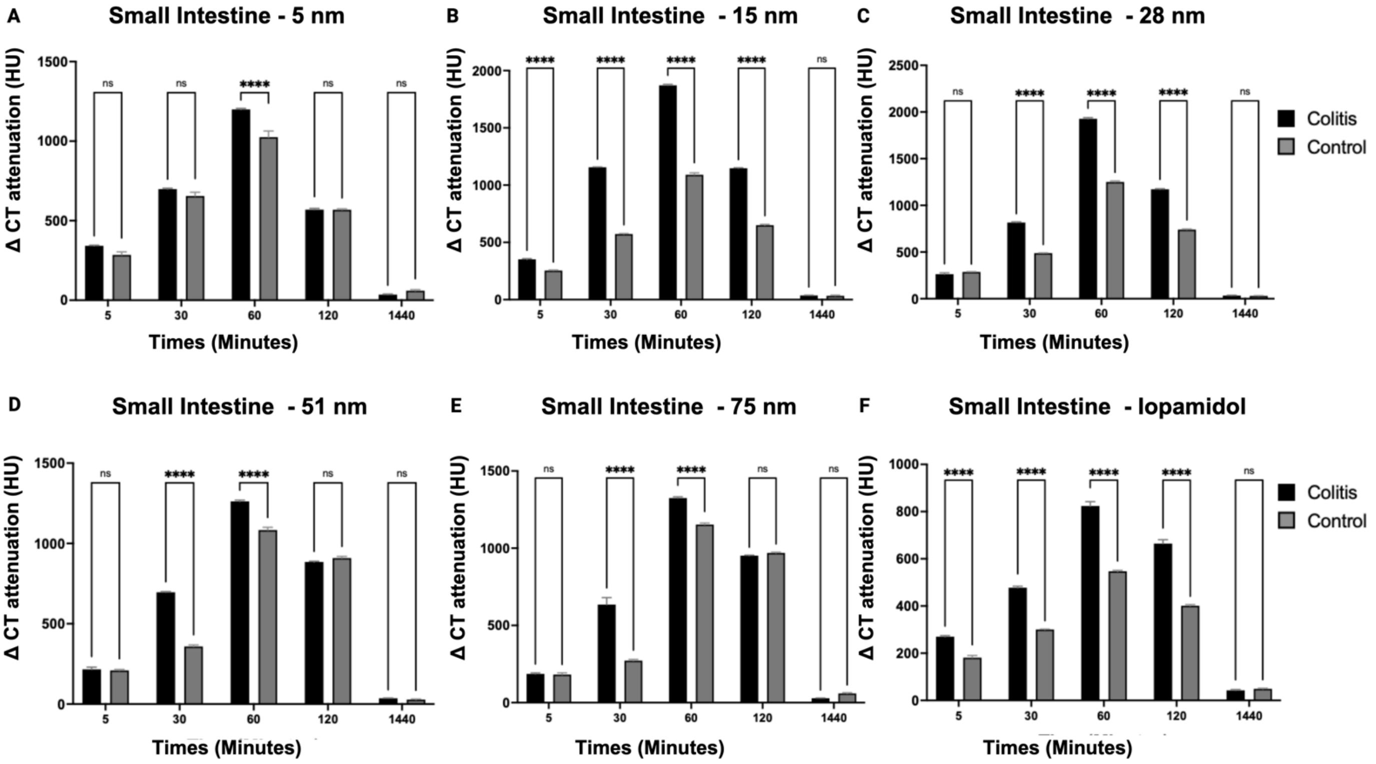


Figure S6: CT attenuation in the small intestine from control and colitis mice at different time points. Error bars represent standard error of the mean, and * Indicates P < 0.05, and **** Indicates P ≤ 0.0001.

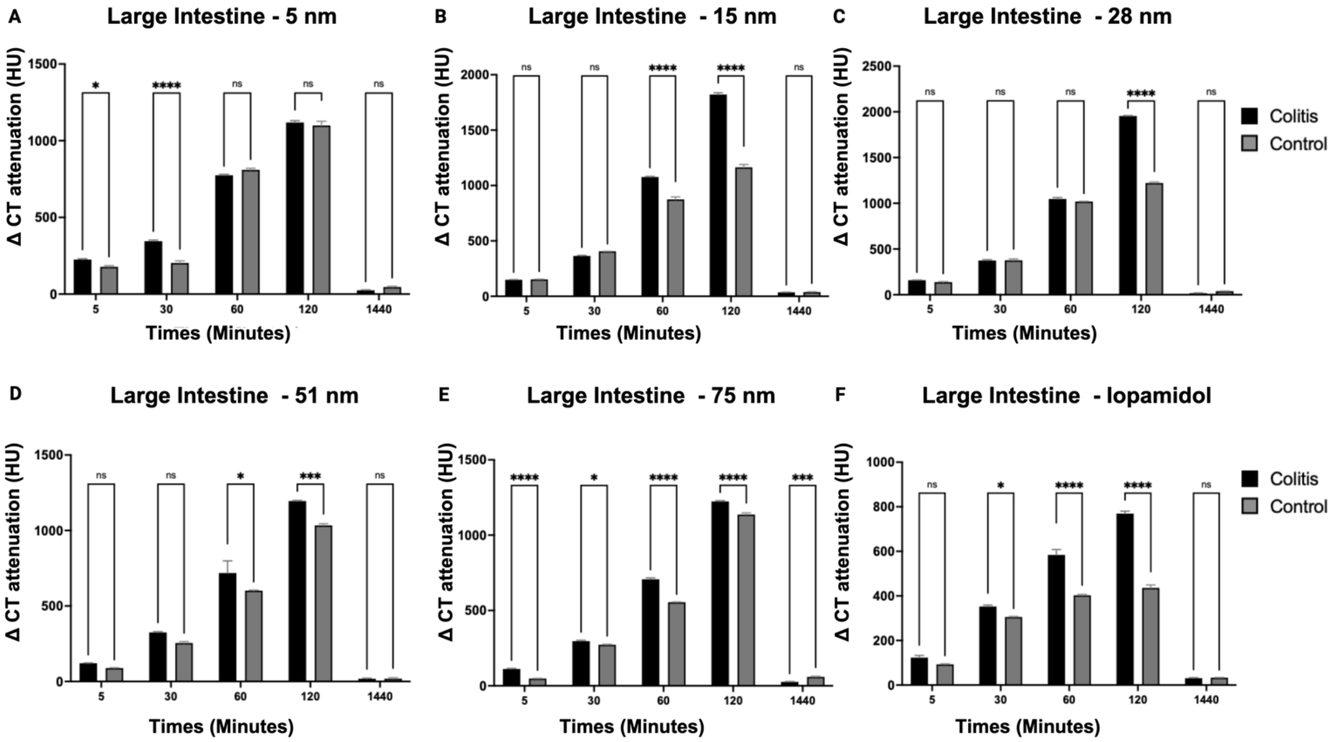


Figure S7: CT attenuation in the large intestine from control and colitis mice at different time points. Error bars represent standard error of the mean, and * Indicates P < 0.05, and **** Indicates P ≤ 0.0001.

| p-value for Stomach - Control group | | | | | |
|---|---|---|---|---|---|
| 5 nm | | 15 nm | | 28 nm | |
| 5 vs. 30 | <0.0001 | 5 vs. 30 | <0.0001 | 5 vs. 30 | 0.0001 |
| 5 vs. 60 | <0.0001 | 5 vs. 60 | <0.0001 | 5 vs. 60 | <0.0001 |
| 5 vs. 120 | <0.0001 | 5 vs. 120 | <0.0001 | 5 vs. 120 | <0.0001 |

| | | | | | |
|---|---|---|---|---|---|
| 5 vs. 1440 | <0.0001 | 5 vs. 1440 | <0.0001 | 5 vs. 1440 | <0.0001 |
| 30 vs. 60 | 0.0007 | 30 vs. 60 | 0.0004 | 30 vs. 60 | <0.0001 |
| 30 vs. 120 | <0.0001 | 30 vs. 120 | <0.0001 | 30 vs. 120 | <0.0001 |
| 30 vs. 1440 | <0.0001 | 30 vs. 1440 | <0.0001 | 30 vs. 1440 | <0.0001 |
| 60 vs. 120 | 0.0010 | 60 vs. 120 | 0.0002 | 60 vs. 120 | <0.0001 |
| 60 vs. 1440 | <0.0001 | 60 vs. 1440 | <0.0001 | 60 vs. 1440 | <0.0001 |
| 120 vs. 1440 | <0.0001 | 120 vs. 1440 | <0.0001 | 120 vs. 1440 | <0.0001 |
| 51 nm | | 75 nm | | Iopamidol | |
| 5 vs. 30 | <0.0001 | 5 vs. 30 | <0.0001 | 5 vs. 30 | <0.0001 |
| 5 vs. 60 | <0.0001 | 5 vs. 60 | <0.0001 | 5 vs. 60 | <0.0001 |
| 5 vs. 120 | <0.0001 | 5 vs. 120 | <0.0001 | 5 vs. 120 | <0.0001 |
| 5 vs. 1440 | <0.0001 | 5 vs. 1440 | <0.0001 | 5 vs. 1440 | <0.0001 |
| 30 vs. 60 | <0.0001 | 30 vs. 60 | <0.0001 | 30 vs. 60 | <0.0001 |
| 30 vs. 120 | <0.0001 | 30 vs. 120 | <0.0001 | 30 vs. 120 | <0.0001 |
| 30 vs. 1440 | <0.0001 | 30 vs. 1440 | <0.0001 | 30 vs. 1440 | <0.0001 |

| | | | | | |
|---|---|---|---|---|---|
| 60 vs. 120 | <0.0001 | 60 vs. 120 | <0.0001 | 60 vs. 120 | 0.0002 |
| 60 vs. 1440 | <0.0001 | 60 vs. 1440 | <0.0001 | 60 vs. 1440 | <0.0001 |
| 120 vs. 1440 | <0.0001 | 120 vs. 1440 | <0.0001 | 120 vs. 1440 | <0.0001 |

Supporting Table S1: Comprehensive p values for CT attenuation changes in the stomach of control group across various time points with different nanoparticles formulation during 24-hour imaging.

| **p-value for Small intestine - Control group** | | | | | |
|---|---|---|---|---|---|
| 5 nm | | 15 nm | | 28 nm | |
| 5 vs. 30 | 0.0005 | 5 vs. 30 | <0.0001 | 5 vs. 30 | <0.0001 |
| 5 vs. 60 | <0.0001 | 5 vs. 60 | <0.0001 | 5 vs. 60 | <0.0001 |
| 5 vs. 120 | 0.0002 | 5 vs. 120 | <0.0001 | 5 vs. 120 | <0.0001 |
| 5 vs. 1440 | 0.0003 | 5 vs. 1440 | <0.0001 | 5 vs. 1440 | <0.0001 |
| 30 vs. 60 | 0.0063 | 30 vs. 60 | <0.0001 | 30 vs. 60 | <0.0001 |
| 30 vs. 120 | 0.1269 | 30 vs. 120 | 0.0003 | 30 vs. 120 | <0.0001 |
| 30 vs. 1440 | <0.0001 | 30 vs. 1440 | <0.0001 | 30 vs. 1440 | <0.0001 |
| 60 vs. 120 | 0.0004 | 60 vs. 120 | <0.0001 | 60 vs. 120 | <0.0001 |

| | | | | | |
|---|---|---|---|---|---|
| 60 vs. 1440 | <0.0001 | 60 vs. 1440 | <0.0001 | 60 vs. 1440 | <0.0001 |
| 120 vs. 1440 | <0.0001 | 120 vs. 1440 | <0.0001 | 120 vs. 1440 | <0.0001 |
| 51 nm | | 75 nm | | Iopamidol | |
| 5 vs. 30 | 0.0002 | 5 vs. 30 | 0.0004 | 5 vs. 30 | 0.0003 |
| 5 vs. 60 | <0.0001 | 5 vs. 60 | <0.0001 | 5 vs. 60 | <0.0001 |
| 5 vs. 120 | <0.0001 | 5 vs. 120 | <0.0001 | 5 vs. 120 | <0.0001 |
| 5 vs. 1440 | <0.0001 | 5 vs. 1440 | 0.0023 | 5 vs. 1440 | 0.0001 |
| 30 vs. 60 | <0.0001 | 30 vs. 60 | <0.0001 | 30 vs. 60 | <0.0001 |
| 30 vs. 120 | <0.0001 | 30 vs. 120 | <0.0001 | 30 vs. 120 | <0.0001 |
| 30 vs. 1440 | <0.0001 | 30 vs. 1440 | <0.0001 | 30 vs. 1440 | <0.0001 |
| 60 vs. 120 | 0.0012 | 60 vs. 120 | <0.0001 | 60 vs. 120 | <0.0001 |
| 60 vs. 1440 | <0.0001 | 60 vs. 1440 | <0.0001 | 60 vs. 1440 | <0.0001 |
| 120 vs. 1440 | <0.0001 | 120 vs. 1440 | <0.0001 | 120 vs. 1440 | <0.0001 |

Supporting Table S2: Comprehensive p values for CT attenuation changes in the small intestine of control group across various time points with different nanoparticles formulation during 24-hour imaging.

| p-value for Large intestine - Control group | | | | | |
|---|---|---|---|---|---|
| 5 nm | | 15 nm | | 28 nm | |
| 5 vs. 30 | 0.2949 | 5 vs. 30 | <0.0001 | 5 vs. 30 | 0.0003 |
| 5 vs. 60 | <0.0001 | 5 vs. 60 | <0.0001 | 5 vs. 60 | <0.0001 |
| 5 vs. 120 | <0.0001 | 5 vs. 120 | <0.0001 | 5 vs. 120 | <0.0001 |
| 5 vs. 1440 | 0.0002 | 5 vs. 1440 | <0.0001 | 5 vs. 1440 | <0.0001 |
| 30 vs. 60 | <0.0001 | 30 vs. 60 | <0.0001 | 30 vs. 60 | <0.0001 |
| 30 vs. 120 | <0.0001 | 30 vs. 120 | <0.0001 | 30 vs. 120 | <0.0001 |
| 30 vs. 1440 | 0.0005 | 30 vs. 1440 | <0.0001 | 30 vs. 1440 | <0.0001 |
| 60 vs. 120 | 0.0003 | 60 vs. 120 | <0.0001 | 60 vs. 120 | <0.0001 |
| 60 vs. 1440 | <0.0001 | 60 vs. 1440 | <0.0001 | 60 vs. 1440 | <0.0001 |
| 120 vs. 1440 | <0.0001 | 120 vs. 1440 | <0.0001 | 120 vs. 1440 | <0.0001 |
| 51 nm | | 75 nm | | Iopamidol | |
| 5 vs. 30 | <0.0001 | 5 vs. 30 | <0.0001 | 5 vs. 30 | <0.0001 |
| 5 vs. 60 | <0.0001 | 5 vs. 60 | <0.0001 | 5 vs. 60 | <0.0001 |
| 5 vs. 120 | <0.0001 | 5 vs. 120 | <0.0001 | 5 vs. 120 | <0.0001 |

| | | | | | |
|---|---|---|---|---|---|
| 5 vs. 1440 | 0.0006 | 5 vs. 1440 | 0.3103 | 5 vs. 1440 | <0.0001 |
| 30 vs. 60 | <0.0001 | 30 vs. 60 | <0.0001 | 30 vs. 60 | <0.0001 |
| 30 vs. 120 | <0.0001 | 30 vs. 120 | <0.0001 | 30 vs. 120 | 0.0014 |
| 30 vs. 1440 | <0.0001 | 30 vs. 1440 | <0.0001 | 30 vs. 1440 | <0.0001 |
| 60 vs. 120 | <0.0001 | 60 vs. 120 | <0.0001 | 60 vs. 120 | 0.1130 |
| 60 vs. 1440 | <0.0001 | 60 vs. 1440 | <0.0001 | 60 vs. 1440 | <0.0001 |
| 120 vs. 1440 | <0.0001 | 120 vs. 1440 | <0.0001 | 120 vs. 1440 | <0.0001 |

Supporting Table S3: Comprehensive p values for CT attenuation changes in the large intestine of control group across various time points with different nanoparticles formulation during 24-hour imaging.

| **p-value for Stomach - Colitis group** | | | | | |
|---|---|---|---|---|---|
| 5 nm | | 15 nm | | 28 nm | |
| 5 vs. 30 | <0.0001 | 5 vs. 30 | <0.0001 | 5 vs. 30 | <0.0001 |
| 5 vs. 60 | <0.0001 | 5 vs. 60 | <0.0001 | 5 vs. 60 | <0.0001 |
| 5 vs. 120 | <0.0001 | 5 vs. 120 | <0.0001 | 5 vs. 120 | <0.0001 |

| | | | | | |
|---|---|---|---|---|---|
| 5 vs. 1440 | <0.0001 | 5 vs. 1440 | <0.0001 | 5 vs. 1440 | <0.0001 |
| 30 vs. 60 | <0.0001 | 30 vs. 60 | <0.0001 | 30 vs. 60 | 0.0002 |
| 30 vs. 120 | <0.0001 | 30 vs. 120 | <0.0001 | 30 vs. 120 | <0.0001 |
| 30 vs. 1440 | <0.0001 | 30 vs. 1440 | <0.0001 | 30 vs. 1440 | <0.0001 |
| 60 vs. 120 | 0.0002 | 60 vs. 120 | <0.0001 | 60 vs. 120 | <0.0001 |
| 60 vs. 1440 | <0.0001 | 60 vs. 1440 | <0.0001 | 60 vs. 1440 | <0.0001 |
| 120 vs. 1440 | <0.0001 | 120 vs. 1440 | <0.0001 | 120 vs. 1440 | <0.0001 |
| 51 nm | | 75 nm | | Iopamidol | |
| 5 vs. 30 | <0.0001 | 5 vs. 30 | <0.0001 | 5 vs. 30 | 0.0058 |
| 5 vs. 60 | <0.0001 | 5 vs. 60 | <0.0001 | 5 vs. 60 | 0.0006 |
| 5 vs. 120 | <0.0001 | 5 vs. 120 | <0.0001 | 5 vs. 120 | <0.0001 |
| 5 vs. 1440 | <0.0001 | 5 vs. 1440 | <0.0001 | 5 vs. 1440 | <0.0001 |
| 30 vs. 60 | <0.0001 | 30 vs. 60 | <0.0001 | 30 vs. 60 | 0.1008 |
| 30 vs. 120 | <0.0001 | 30 vs. 120 | <0.0001 | 30 vs. 120 | 0.0043 |
| 30 vs. 1440 | <0.0001 | 30 vs. 1440 | <0.0001 | 30 vs. 1440 | <0.0001 |

| | | | | | |
|---|---|---|---|---|---|
| 60 vs. 120 | <0.0001 | 60 vs. 120 | <0.0001 | 60 vs. 120 | 0.1481 |
| 60 vs. 1440 | <0.0001 | 60 vs. 1440 | <0.0001 | 60 vs. 1440 | 0.0001 |
| 120 vs. 1440 | <0.0001 | 120 vs. 1440 | <0.0001 | 120 vs. 1440 | <0.0001 |

Supporting Table S4: Comprehensive p values for CT attenuation changes in the stomach across various time points in colitis mice treated with different nanoparticles formulation during 24-hour imaging.

| **p-value for Small intestine - Colitis group** | | | | | |
|---|---|---|---|---|---|
| 5 nm | | 15 nm | | 28 nm | |
| 5 vs. 30 | <0.0001 | 5 vs. 30 | <0.0001 | 5 vs. 30 | <0.0001 |
| 5 vs. 60 | <0.0001 | 5 vs. 60 | <0.0001 | 5 vs. 60 | <0.0001 |
| 5 vs. 120 | <0.0001 | 5 vs. 120 | <0.0001 | 5 vs. 120 | <0.0001 |
| 5 vs. 1440 | <0.0001 | 5 vs. 1440 | <0.0001 | 5 vs. 1440 | <0.0001 |
| 30 vs. 60 | <0.0001 | 30 vs. 60 | <0.0001 | 30 vs. 60 | <0.0001 |
| 30 vs. 120 | <0.0001 | 30 vs. 120 | 0.4742 | 30 vs. 120 | <0.0001 |
| 30 vs. 1440 | <0.0001 | 30 vs. 1440 | <0.0001 | 30 vs. 1440 | <0.0001 |
| 60 vs. 120 | <0.0001 | 60 vs. 120 | <0.0001 | 60 vs. 120 | <0.0001 |

| | | | | | |
|---|---|---|---|---|---|
| 60 vs. 1440 | <0.0001 | 60 vs. 1440 | <0.0001 | 60 vs. 1440 | <0.0001 |
| 120 vs. 1440 | <0.0001 | 120 vs. 1440 | <0.0001 | 120 vs. 1440 | <0.0001 |
| 51 nm | | 75 nm | | Iopamidol | |
| 5 vs. 30 | <0.0001 | 5 vs. 30 | 0.0013 | 5 vs. 30 | <0.0001 |
| 5 vs. 60 | <0.0001 | 5 vs. 60 | <0.0001 | 5 vs. 60 | <0.0001 |
| 5 vs. 120 | <0.0001 | 5 vs. 120 | <0.0001 | 5 vs. 120 | <0.0001 |
| 5 vs. 1440 | 0.0003 | 5 vs. 1440 | <0.0001 | 5 vs. 1440 | <0.0001 |
| 30 vs. 60 | <0.0001 | 30 vs. 60 | <0.0001 | 30 vs. 60 | <0.0001 |
| 30 vs. 120 | <0.0001 | 30 vs. 120 | 0.0050 | 30 vs. 120 | 0.0002 |
| 30 vs. 1440 | <0.0001 | 30 vs. 1440 | 0.0002 | 30 vs. 1440 | <0.0001 |
| 60 vs. 120 | <0.0001 | 60 vs. 120 | <0.0001 | 60 vs. 120 | <0.0001 |
| 60 vs. 1440 | <0.0001 | 60 vs. 1440 | <0.0001 | 60 vs. 1440 | <0.0001 |
| 120 vs. 1440 | <0.0001 | 120 vs. 1440 | <0.0001 | 120 vs. 1440 | <0.0001 |

Supporting Table S5: Comprehensive p values for CT attenuation changes in the small intestine across various time points in colitis mice treated with different nanoparticles formulation during 24-hour imaging.

| p-value for Large intestine - Colitis group | | | | | |
|---|---|---|---|---|---|
| 5 nm | | 15 nm | | 28 nm | |
| 5 vs. 30 | 0.0002 | 5 vs. 30 | <0.0001 | 5 vs. 30 | <0.0001 |
| 5 vs. 60 | <0.0001 | 5 vs. 60 | <0.0001 | 5 vs. 60 | <0.0001 |
| 5 vs. 120 | <0.0001 | 5 vs. 120 | <0.0001 | 5 vs. 120 | <0.0001 |
| 5 vs. 1440 | <0.0001 | 5 vs. 1440 | <0.0001 | 5 vs. 1440 | <0.0001 |
| 30 vs. 60 | <0.0001 | 30 vs. 60 | <0.0001 | 30 vs. 60 | <0.0001 |
| 30 vs. 120 | <0.0001 | 30 vs. 120 | <0.0001 | 30 vs. 120 | <0.0001 |
| 30 vs. 1440 | <0.0001 | 30 vs. 1440 | <0.0001 | 30 vs. 1440 | <0.0001 |
| 60 vs. 120 | <0.0001 | 60 vs. 120 | <0.0001 | 60 vs. 120 | <0.0001 |
| 60 vs. 1440 | <0.0001 | 60 vs. 1440 | <0.0001 | 60 vs. 1440 | <0.0001 |
| 120 vs. 1440 | <0.0001 | 120 vs. 1440 | <0.0001 | 120 vs. 1440 | <0.0001 |
| 51 nm | | 75 nm | | Iopamidol | |
| 5 vs. 30 | <0.0001 | 5 vs. 30 | <0.0001 | 5 vs. 30 | <0.0001 |
| 5 vs. 60 | 0.0035 | 5 vs. 60 | <0.0001 | 5 vs. 60 | <0.0001 |
| 5 vs. 120 | <0.0001 | 5 vs. 120 | <0.0001 | 5 vs. 120 | <0.0001 |

| | | | | | |
|---|---|---|---|---|---|
| 5 vs. 1440 | <0.0001 | 5 vs. 1440 | <0.0001 | 5 vs. 1440 | 0.0012 |
| 30 vs. 60 | 0.0199 | 30 vs. 60 | <0.0001 | 30 vs. 60 | 0.0016 |
| 30 vs. 120 | <0.0001 | 30 vs. 120 | <0.0001 | 30 vs. 120 | <0.0001 |
| 30 vs. 1440 | <0.0001 | 30 vs. 1440 | <0.0001 | 30 vs. 1440 | <0.0001 |
| 60 vs. 120 | 0.0092 | 60 vs. 120 | <0.0001 | 60 vs. 120 | 0.0022 |
| 60 vs. 1440 | 0.0016 | 60 vs. 1440 | <0.0001 | 60 vs. 1440 | <0.0001 |
| 120 vs. 1440 | <0.0001 | 120 vs. 1440 | <0.0001 | 120 vs. 1440 | <0.0001 |

Supporting Table S6: Comprehensive p values for CT attenuation changes in the large intestine across various time points in colitis mice treated with different nanoparticles formulation during 24-hour imaging.

| **p-value for control group** | | | | | |
|---|---|---|---|---|---|
| Stomach – 5 minutes Time point | | Small intestine– 60 minutes Time point | | Large intestine– 120 minutes Time point | |
| 5 vs. 15 | 0.9922 | 5 vs. 15 | 0.1886 | 5 vs. 15 | 0.1555 |
| 5 vs. 25 | <0.0001 | 5 vs. 25 | <0.0001 | 5 vs. 25 | 0.0006 |
| 5 vs. 50 | <0.0001 | 5 vs. 50 | 0.3028 | 5 vs. 50 | 0.1373 |

| | | | | | |
|---|---|---|---|---|---|
| 5 vs. 75 | <0.0001 | 5 vs. 75 | 0.0008 | 5 vs. 75 | 0.7089 |
| 5 vs. ISO | <0.0001 | 5 vs. ISO | <0.0001 | 5 vs. ISO | <0.0001 |
| 15 vs. 25 | <0.0001 | 15 vs. 25 | <0.0001 | 15 vs. 25 | 0.2371 |
| 15 vs. 50 | <0.0001 | 15 vs. 50 | 0.9998 | 15 vs. 50 | 0.0003 |
| 15 vs. 75 | <0.0001 | 15 vs. 75 | 0.2396 | 15 vs. 75 | 0.8886 |
| 15 vs. ISO | <0.0001 | 15 vs. ISO | <0.0001 | 15 vs. ISO | <0.0001 |
| 25 vs. 50 | 0.1038 | 25 vs. 50 | <0.0001 | 25 vs. 50 | <0.0001 |
| 25 vs. 75 | 0.0037 | 25 vs. 75 | 0.0138 | 25 vs. 75 | 0.0247 |
| 25 vs. ISO | <0.0001 | 25 vs. ISO | <0.0001 | 25 vs. ISO | <0.0001 |
| 50 vs. 75 | 0.7342 | 50 vs. 75 | 0.1443 | 50 vs. 75 | 0.0047 |
| 50 vs. ISO | <0.0001 | 50 vs. ISO | <0.0001 | 50 vs. ISO | <0.0001 |
| 75 vs. ISO | <0.0001 | 75 vs. ISO | <0.0001 | 75 vs. ISO | <0.0001 |

Supporting Table S7: P Values for Attenuation Dynamics of AuNP (5-75 nm) and Iopamidol in the gastrointestinal tract of control group mice across different organs and time points.

| **p-value for colitis group** |
|---|

| Stomach – 5 minutes Time point | | Small intestine– 60 minutes Time point | | Large intestine– 120 minutes Time point | |
|---|---|---|---|---|---|
| 5 vs. 15 | <0.0001 | 5 vs. 15 | <0.0001 | 5 vs. 15 | <0.0001 |
| 5 vs. 25 | <0.0001 | 5 vs. 25 | <0.0001 | 5 vs. 25 | <0.0001 |
| 5 vs. 50 | <0.0001 | 5 vs. 50 | 0.0050 | 5 vs. 50 | 0.0001 |
| 5 vs. 75 | <0.0001 | 5 vs. 75 | <0.0001 | 5 vs. 75 | <0.0001 |
| 5 vs. ISO | <0.0001 | 5 vs. ISO | <0.0001 | 5 vs. ISO | <0.0001 |
| 15 vs. 25 | 0.0003 | 15 vs. 25 | 0.0151 | 15 vs. 25 | <0.0001 |
| 15 vs. 50 | <0.0001 | 15 vs. 50 | <0.0001 | 15 vs. 50 | <0.0001 |
| 15 vs. 75 | 0.0158 | 15 vs. 75 | <0.0001 | 15 vs. 75 | <0.0001 |
| 15 vs. ISO | <0.0001 | 15 vs. ISO | <0.0001 | 15 vs. ISO | <0.0001 |
| 25 vs. 50 | 0.1266 | 25 vs. 50 | <0.0001 | 25 vs. 50 | <0.0001 |
| 25 vs. 75 | 0.6447 | 25 vs. 75 | <0.0001 | 25 vs. 75 | <0.0001 |
| 25 vs. ISO | <0.0001 | 25 vs. ISO | <0.0001 | 25 vs. ISO | <0.0001 |
| 50 vs. 75 | 0.0032 | 50 vs. 75 | 0.0049 | 50 vs. 75 | 0.3565 |

| | | | | | |
|---|---|---|---|---|---|
| 50 vs. ISO | <0.0001 | 50 vs. ISO | <0.0001 | 50 vs. ISO | <0.0001 |
| 75 vs. ISO | <0.0001 | 75 vs. ISO | <0.0001 | 75 vs. ISO | <0.0001 |

Supporting Table S8: P Values for Attenuation Dynamics of AuNP (5-75 nm) and Iopamidol in the gastrointestinal tract of colitis group mice across different organs and time points.

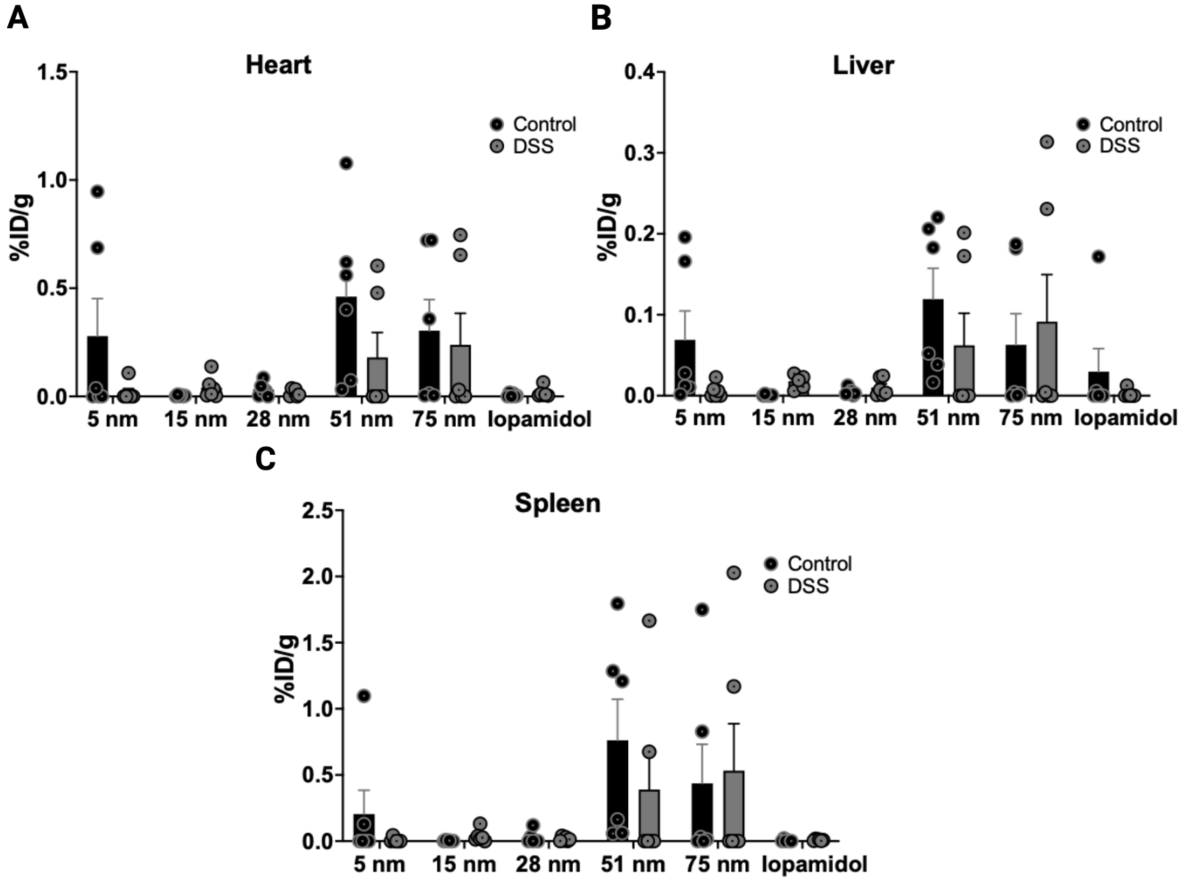


Figure S8: Biodistribution of AuNP in healthy and DSS-colitis mice at 24 hrs post administration.
